# Motile cilia spin the Reissner fiber, a tensioned and anchored extracellular thread essential for body morphogenesis

**DOI:** 10.1101/2025.09.25.678623

**Authors:** Elizabeth A. Bearce, Samuel G. Bertrand, Samara Williams, Sophie I. Fisher, Adamend L. Freda, Zoe H. Irons, Calvin Chmelir, Daniel T. Grimes

## Abstract

The Reissner fiber (RF) is an extracellular thread composed of the glycoprotein SCO-spondin (Sspo) that forms within cerebrospinal fluid (CSF) in the brain ventricles and central canal (CC) of vertebrates. Its assembly depends on motile cilia, yet how cilia and CSF flow transform secreted Sspo into a fiber that spans the length of the body axis has remained unknown. Using live imaging in zebrafish, we show that globular Sspo is remodeled into thin fibrils that anchor to the floor plate (FP), elongate, and spin together into a tensioned, posteriorly translocating RF. Fiber assembly draws on a shared CSF pool of Sspo but requires local ciliary activity for globule-to-fibril remodeling. Once assembled, the RF is anchored by dynamic fibrils that stabilize the fiber and preserve its tension. Perturbations that locally disrupt ciliary motility and RF integrity along the CC result in region-specific defects in axial morphogenesis. These findings identify the RF as a flow-responsive extracellular system that converts ciliary beating into mechanical structure, linking fluid dynamics to the large-scale shaping of the vertebrate body.

**Highlights:**

- Motile cilia remodel Sspo globules into FP-anchored fibrils that bundle into RF.
- RF assembly draws on a shared CSF Sspo pool but requires local cilia activity.
- Laser ablation reveals patterned RF tension constrained by dynamic FP anchors.
- Regional cilia defects cause local RF loss and corresponding body shape defects.

## Introduction

Fluid-filled tubes are a widespread feature of multicellular life, including vertebrate neural canals, vasculature, and kidney nephrons, as well as the xylem and phloem of plants. The flows within such tubes serve diverse functions, from transporting nutrients and wastes to dispersing chemical signals, and even constructing extracellular assemblies such as the otoliths of the inner ear (Riley et al. (1997); Freund et al. (2012)). Flows influence embryonic patterning (Nonaka et al. (1998); Sawamoto et al. (2006)), morphogenesis (Chow et al. (2022); Serluca et al. (2002)), host-microbe interactions (Nawroth et al. (2017)) and cancer cell dissemination (Follain et al. (2018)). Despite these roles, the mechanisms by which flows remodel the extracellular environment to influence development and physiology remain elusive.

First described in 1860, the Reissner fiber (RF) is a continuous, polymeric thread that forms in the cerebrospinal fluid (CSF) of the brain ventricles and central canal (CC) of the spinal cord (Reissner (1860); Sepúlveda et al. (2021); Aboitiz and Montiel (2021)). The RF is composed primarily of SCO-spondin (Sspo), a >5,000-amino acid glycoprotein secreted during early development by the floor plate (FP) at the base of the neural tube and, later, by secretory circumventricular organs of the brain (Rodríguez et al. (1998); Yulis et al. (1998); Sepúlveda et al. (2021)). In zebrafish larvae, motile cilia on ependymal precursor cells line the CC, generating both local flow vortices and bulk movements of CSF (Kramer-Zucker et al. (2005); Grimes et al. (2016); Thouvenin et al. (2020)). These flows are required for RF assembly: when ciliary beating is absent and CSF circulation is disrupted, Sspo is still secreted into the CC but remains diffuse and flocculent rather than organized into a fiber (Cantaut-Belarif et al. (2018); Troutwine et al. (2020); Bearce et al. (2022b)). How motile cilia remodel Sspo into the RF, however, remains an open question.

Once assembled, the RF lies under tension within the CC and translocates caudally at a rate that varies across chordates (Dendy (1909); Troutwine et al. (2020); Bellegarda et al. (2023)). Renewal of the entire thread occurs variably, with turnover estimated at 3-4 days in frogs, 6-8 days in zebrafish, and 10-40 days in mice and goldfish (Ermisch et al. (1970); Diederen (1973); Sterba et al. (1967); Leatherland and Dodd (1968)). The mechanisms generating tension and driving translocation remain unclear, though motile cilia are likely contributors: during beating, they directly contact the RF (Troutwine et al. (2020)), suggesting their forces could sustain tension and propel fiber movement.

The developmental roles of the RF are also coming into focus. Anatomists long speculated that the RF might influence body shape (Dendy (1909); Nicholls (1913)), and subsequent resection studies lent experimental support (Hauser (1972); Rühle (1971)). More recently, genetic analyses have confirmed that disrupting RF formation leads to defects in axial morphology (Cantaut-Belarif et al. (2018); Lu et al. (2020); Troutwine et al. (2020); Bearce et al. (2022b); Wang et al. (2022); Xu et al. (2023); Rose et al. (2020)). In zebrafish, shortly after CC opening, the embryo undergoes axial straightening morphogenesis in which the initially ventrally curved embryo straightens its body axis (Kimmel et al. (1995); Bearce and Grimes (2020)). When the RF is absent—either in cilia motility mutants that fail to assemble it from secreted Sspo, or in mutants lacking Sspo altogether—straightening fails and embryos retain a ventral curve (Kramer-Zucker et al. (2005); Cantaut-Belarif et al. (2018); Bearce and Grimes (2020)). Moreover, hypomorphic sspo mutants that initially straighten later develop scoliosis-like curves (Rose et al. (2020); Troutwine et al. (2020)). Thus, the RF plays functional roles in establishing the linear body axis during embryonic stages and maintaining spinal alignment during growth.

Here, using live imaging of RF assembly from secreted Sspo in zebrafish, we show that globules of Sspo are drawn into thin fibrils that anchor to the FP, elongate, twist together, and bundle into a continuous RF. This transformation resembles the spinning of wool, in which loose, flocculent fibers are drawn and twisted into a laterally stabilized thread. Genetic mosaics reveal RF assembly draws on a common CSF pool of Sspo but requires local motile cilia activity for fiber growth and maintenance. Photoablation demonstrates that the mature RF exists under spatially patterned tension constrained by fibrillar anchors linking it to the FP. Finally, analysis of mutants with spatially or temporally reduced cilia motility shows that RF assembly tightly correlates with regional body straightening, suggesting local morphogenetic consequences of the RF. Together, these results explain how motile cilia remodel secreted Sspo into a mechanically anchored, flow-responsive fiber and connect its assembly to vertebrate body shaping.

## Results

### RF assembly requires motile cilia-driven remodeling of secreted Sspo

#### Stepwise RF assembly

To determine how the RF assembles, we combined differential interference contrast imaging to visualize motile cilia with live and fixed fluorescence imaging of an Sspo-GFP knock-in line (Troutwine et al. (2020)). This revealed four phases of RF formation, schematized in Fig 1A.

**Figure 1.**
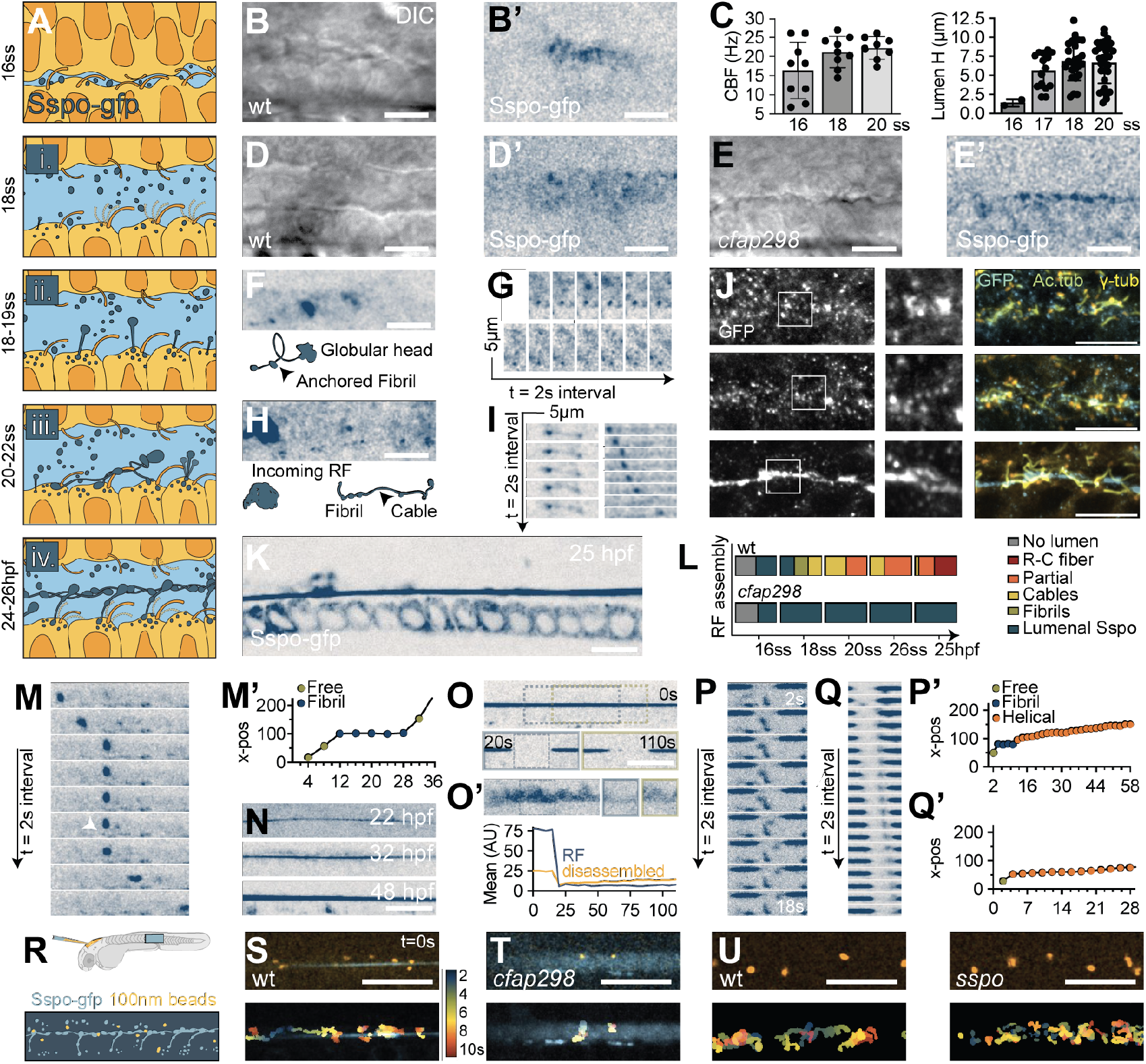
RF assembly. **A)** Schematic phases of RF assembly. **B-B’)** Early lumen opening (B) and Sspo-GFP secretion (B’) at 16 ss. **C)** Median cilia beat frequency (CBF) and lumen height during 16-20 ss. Error bars show s.d. **D-D’)** By 18 ss, the CC lumen is contiguous (D) and contains flocculent Sspo-GFP (D’). **E-E’)** *cfap298* mutants exhibit narrow CC lumina (E) with dense Sspo-GFP accumulation (E’). **F-G)** Sspo-GFP extrudes into fibrils (F) from stably anchored membrane puncta that persist over time (G) and are capped by a globular head. **H)** Fibrils from neighboring FP cells elongate and join to form cables (arrowheads) **I)** Cables remain membrane-associated and mediate posterior transfer of Sspo-GFP material. **J)** Immunostaining for GFP (Sspo-GFP), acetylated *α*-tubulin (cilia) and *γ*-tubulin (microtubule organizing centers) at 16-20 ss. **K)** RF assembly is contiguous along the CC by 25 hpf. **L)** Qualitative phenotyping of RF assembly across developmental stages. **M-M’)** RF material dislodged from an anterior CC region is caught by a membrane-bound fibril (M, arrowhead), which transiently slows its caudal progression down the CC (M’). **N)** The RF matures over time, incorporating additional material. **O-O’)** 10 µm regions (dashed boxes) of 28 hpf RF were photobleached (t = 20 s) and show little fluorescence recover over minutes (t = 110 s). Secreted disorganized material instead from *cfap298* mutants shows fluorescence recovery (O’). The graph quantifies fluorescence changes in indicated regions. **P-P’)** Free-floating Sspo is captured by a fibril before helical deposition along a photobleached segment of the RF. Timelapse shows data are given (P) and particle trajectory quantified (P’). **Q-Q’)** Free-floating Sspo is deposited within a photobleached segment of the RF (Q) and translocates caudally (Q’) during helical rotation. **R)** Schematic of ventricle bead injections at 28 hpf, several hours prior to CC imaging. **S-U’)** Imaging of Sspo-GFP and beads in WT (S), *cfap298* (T), *sspo* heterozygotes (U) and *sspo* mutants (U’). Temporal hyperstack (lower) demonstrates laminar and helical flow trajectories close to the RF in WT and *sspo* heterozygotes (S, U). Reduced cilia motility prevents RF assembly and beads demonstrate trajectories with low displacement (T). Increased and erratic mixing was observed in *sspo* mutants (U’). Scale bars: 10 µm.

Stage 1 began at the 16-somite stage (ss), when the central canal (CC) lumen opened as small, discontinuous fluid-filled pockets along the midline (Fig 1A-B and Video 1). Each pocket contained a beating cilium, suggesting that nascent ciliary motility contributed to CC opening (Video 1). Between 16-18 ss, ciliary motility increased, and pockets widened and coalesced to generate a contiguous CC lumen by 18 ss (Kimmel et al. (1995))(Fig 1Ai, 1C-D and Video 2). Concurrently, Sspo secreted from floor plate (FP) cells accumulated as globular material within the lumen (Fig 1B’, 1D’ and Video 3).

In Stage 2 (18-19 ss), luminal Sspo exhibited two behaviors: some globules remained at the FP apical surface, while others moved within the nascent cerebrospinal fluid (CSF) (Video 4). Next, in the vicinity of a motile cilium, FP-associated globules appeared to be captured in local flows and changed conformation, being remodeled from a globular shape to a thin fibrillar shape that we term fibrils. These fibrils maintained an attachment point at the FP membrane and contained a globular head that extended into the luminal space of the CC (Fig 1Aii, 1D’, 1F-G and Video 5).

In Stage 3 (20-22 ss), additional secreted Sspo streamed along fibrils, a process which lengthened fibrils and increased the size of their globular heads (Fig 1G and Video 6). Once adjacent fibrils from neighboring FP cells extended far enough to contact one another, they formed lateral associations that twisted the fibrils together (Fig 1Aiii and 1H-1J). This generated discontinuous patches of FP cells throughout the CC that were linked by thin streams of fibrillar Sspo. We refer to these linked fibrils as cables. Cables displayed caudally moving, bead-like clusters of Sspo material along their length, likely reflecting continued incorporation of Sspo (Fig 1I and Video 7).

In Stage 4 (22 ss-26 hpf), cable-connected FP cells appeared to increase Sspo output, reinforcing connections and thickening the forming RF, based on increased Sspo-GFP fluorescence intensity. Globular aggregates continued to move along cables, and these bundled tracts slowly extended along the CC. This process ultimately yielded a mature, continuous RF throughout the rostro-caudal extent of the CC by 25 hpf in most individuals (Fig 1Aiv, 1K-L and Video 8). Super-resolution imaging of early fibrils revealed a beads-on-a-string appearance, which likely reflects a higher-order repeating of Sspo-GFP subunits within the fibril (Fig 1J).

#### Instability and refinement

Although RF formation proceeded in this stepwise manner along the entire central canal, where FP-anchored fibrils in no particular anterior-posterior position were competent to nucleate short stretches of RF assembly, immature fibers were labile and prone to breakage. Stretches of fibrillar material occasionally disassembled or detached from the FP and were advected rapidly toward the caudal CC (Fig 1M-M’ and Video 9). During transit, detached material did not retain its fibrillar conformation but instead collapsed into bright globular masses while moving down the CC. These segments then transiently captured on FP-bound fibrils (Fig 1M’ and Video 9). These breakdown events coincided with accumulation of Sspo material in the caudal CC, suggesting that early instability and turnover contributed to posterior Sspo enrichment (Video 10). Eventually a highly stabilized RF is nucleated at a far anterior region of the CC which exhibits a more rapid A-P assembly along early fibril and cable sites (Fig 1H). Thus, RF assembly involved repeated cycles of formation and refinement. This likely reflects a requirement for sufficient Sspo material accumulation in the CC and multiple FP attachment sites before a continuous fiber can be maintained.

#### Loss of fibril remodeling in motility-deficient conditions

Ciliary motility in the CC is essential for RF formation (Cantaut-Belarif et al. (2018)). Our time-course analysis suggested motile cilia contribute to multiple stages of the RF assembly process.

To test this directly, we used temperature-sensitive *cfap298* mutants, which display strongly reduced, but not absent, ciliary motility when raised at 28°C (Jaffe et al. (2016); Bearce et al. (2022a)). In mutants, although a contiguous CC lumen does form, its opening was delayed and the lumen was narrower and irregular in shape, exhibiting a wavy morphology (Fig 1E). Importantly, when imaging Sspo-GFP, we did not observe a transition to fibrillar material (Stage 2) in *cfap298* mutants (Fig 1E’ and 1L). Sspo continued to be secreted from the FP in mutants but, rather than forming a fiber, instead accumulated in the CC in a globular state (Fig 1E’ and Video 11). This indicates that ciliary motility is required for remodeling Sspo into fibrils and thus for initiating RF scaffold formation. Since the globule-to-fibril transition consistently occurred at apical sites near motile cilia in wild type and was lost in motility-deficient mutants, we suggest ciliary beating provides mechanical forces that drive this conformational change.

#### Material addition during RF growth

After initially forming as a thin fiber, the RF increases in thickness as development progresses (Fig 1N). To assess how secreted material incorporated into the developing fiber to promote this growth, we photobleached short RF segments and tracked new incorporation. The assembled RF was stable over minutes, showing minimal fluorescence recovery after photobleaching (FRAP)(Fig 1O), in agreement with prior work (Troutwine et al. (2020); Bellegarda et al. (2023)). By comparison, unassembled Sspo in the CC of *cfap298* mutants showed fluorescent recovery, indicating higher free exchange of bleached Sspo-GFP (Fig 1O’). Despite the high stability of assembled RF, we did observe sparse incorporation of new material along bleached regions from flocculent particles of Sspo within the CC. During this incorporation, globular Sspo undergoing rapid vortical motion in the CSF transiently caught on FP-attached fibrils, leading to reduced movement, and was then passed along fibrils into the RF (Fig 1P-P’ and Video 12). As they incorporated, globules spun around the RF and then began to move away from the fibril, translocating caudally at rates of 100-300 nm s^*−*^1, consistent with the rate of RF translocation down the CC (Fig 1Q-Q’ and Video 12). These observations indicate that fiber growth is sustained by continuous recruitment of secreted material. Sspo addition during RF growth proceeds similarly to initial fibril formation, with globular Sspo being mechanically extruded by local shear at the site of FP-attachment and then wrapped helically around the RF to support its growth in a mechanism that is analogous to the spinning of cotton candy.

#### Hydrodynamics in the CC are altered by the RF

Because Sspo material moved helically around the RF during incorporation, we next assayed CSF dynamics to determine whether this was a consequence of fluid flow. By injecting 100 nm fluorescent microspheres (beads) into the brain ventricles, we tracked bead movement as they transited along the CC (Fig 1R). Bead movement progressed rostral to caudal over several hours with a visible front. In wild type, individual beads often followed helical trajectories within local vortices, including spinning around the RF (Fig 1S and Video 13), consistent with the helical motion of incorporating Sspo. When near the RF, beads occasionally shifted rapidly along the fiber, in either rostral or caudal directions, transitioning between adjacent vortical domains (Video 14). This was dependent on motile cilia, as bead transit rates were significantly reduced in the hypofunctional ciliary motility environment of *cfap298* mutants, where movement was reduced, and tracks were vortical but with low longitudinal displacement (Fig 1T). This suggests that the RF can reshape local flow streamlines and may provide a conduit to help particles escape otherwise stable recirculation zones, thereby enhancing longitudinal transport of material in the CC.

Unexpectedly, in *sspo* mutants that lacked Sspo secretion and RF assembly, rostral-to-caudal bead mixing within the CC was increased compared to wild type (Fig 1U and Video 15). One possibility is altered CSF rheology: concentrated Sspo, a large glycoprotein, could increase the effective viscosity of CSF, leading to less effective cilia-driven mixing. The absence of Sspo, leading to a less viscous CSF, might then cause increased mixing.

#### Summary

In summary, we find that RF assembly proceeds through four stages (Fig 1A): (1) CC opening and Sspo secretion; (2) a globular-to-fibril transition of Sspo material near motile cilia; (3) fibril elongation and lateral association into cables; and (4) continued Sspo addition and posterior transport, leading to RF thickening. Early extensions are labile and prone to breakage before stabilization across multiple FP attachments. Motile cilia are required to initiate fibril formation and likely support later steps by promoting elongation and encounters between fibrils. During fiber growth and maintenance, globular Sspo was caught on fibrils and then spun into the main fiber. Finally, Sspo and the RF alter hydrodynamics within the CC, constraining cilia-driven mixing and potentially guiding long-range particle transport.

### Local cilia motility is required for RF assembly, whereas Sspo acts non-locally

We next tested whether RF assembly depends on local Sspo secretion or a distributed source within the CC. We considered two models: (1) RF assembly requires Sspo secretion from FP cells directly beneath the forming fiber, or (2) Sspo released anywhere along the CC enters a shared CSF pool and contibutes to RF formation regardless of origin (Fig 2A). Our imaging in wild type was consistent with both models: we observed fibrils on FP cells incorporating Sspo material into the RF locally, supporting local secretion, but we also documented free-floating Sspo globules that elongated and integrated into the fiber upon contacting motile cilia, favoring distributed assembly.

**Figure 2.**
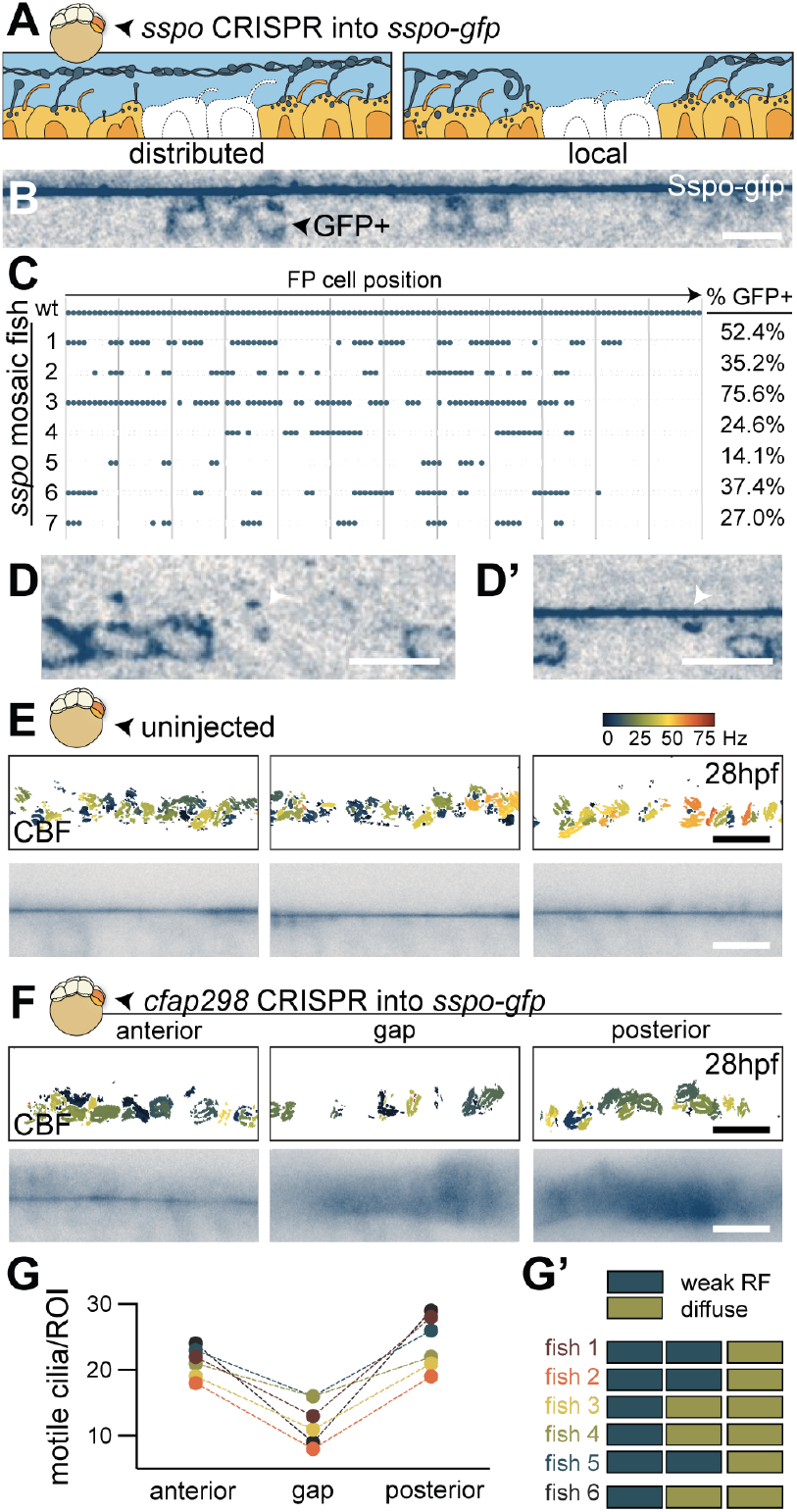
Mosaic analysis. **A)** Schematic of distributed and local mechanisms of RF assembly which were tested by injecting sspo CRISPR into 1 cell at the 8-cell stage to generate mosaic embryos. **B)** RF assembly in a 28 hpf embryo with mosaic expression of FP Sspo-GFP. **C)** Spatial distribution of GFP+ FP cells in the caudal end of 7 embryos with intact RF and mosaic Sspo-GFP depletion. Circles represent GFP+ cells. **D-D’)** Examples of fibrils attached to GFP-FP cells, with (D’) or without (D) overlying RF. **E-F)** Heat map of CBF (top) and Sspo-GFP (bottom) of control (E) or *cfap298* mosaics (F) in regions surrounding gaps in robust cilia motility and equivalent regions in controls. **G-G’)** Quantitation of cilia motility and RF integrity in *cfap298* mosaic fish. The number of motile cilia was scored within regions of interest (ROIs) at motility-deficient gaps and adjacent anterior and posterior regions in 6 mosaic fish (G). In the same individuals, Sspo-GFP signal was assessed as either weak RF or diffuse (G’). Scale bars: 10 µm.

To test these models genetically, we generated *sspo* mosaic embryos by injecting Cas9 and *sspo*-targeting gRNAs into homozygous Sspo-GFP embryos, producing a patchy distribution of GFP+ (Sspo-expressing) and GFP-(Sspo mutant) FP cells. This preserved CC architecture while varying local Sspo availability. At high gRNA doses (1000 pg per embryo), mutagenesis was widespread, and RF formation failed. At lower doses (10 pg per embryo), many FP cells retained Sspo-GFP expression, and the RF was able to assemble in the CC (Fig 2B-C). Notably, RF formation spanned across regions where several adjacent FP cells lacked Sspo-GFP expression (Fig 2B-C). This supports the pooled secretion model: Sspo can travel through the CSF and contribute to RF assembly at sites distant from its release. Indeed, we found instances of GFP-cells that harbored a surface Sspo fibril, suggesting that fibril had formed from Sspo secreted elsewhere in the CC (Fig 2D-D’ and Video 16). Moreover, these data demonstrate that RF assembly is robust, tolerating local gaps in secretion and being able to self-organize from a distributed Sspo source.

We next asked whether cilia motility is locally required for RF assembly by generating *cfap298* mosaics with discrete motility-deficient patches along the CC. In each embryo, we imaged the motility-deficient region (termed the ‘gap’) together with immediately adjacent rostral and caudal regions (Fig 2E-F). We found that RF segments formed rostral to the motility-deficient zone. In addition to RF, those regions accumulated excess flocculent Sspo, consistent with inefficient incorporation into the fiber under reduced motility (Fig 2F-G’). However, just after entering the motility-deficient zone, the RF consistently broke down and did not reform caudally (Fig 2F-G’). This indicated that although rostral cilia could initiate RF growth, continued elongation and stability of the fiber required local motility, supporting the idea that cilia generate the local hydrodynamic environment for RF assembly and extension down the CC.

Together, these findings suggest that while Sspo secretion near motile cilia may enhance fibril elongation and incorporation of Sspo material into the RF, it is not strictly required at any given axial location. Free Sspo globules can circulate until they encounter local hydrodynamic environments that favor their remodeling into conformations allowing integration into the RF. Thus, RF assembly draws on both a shared molecular Sspo pool and localized remodeling by cilia.

### Ciliary motility maintains RF stability and limits its translocation

Next, we investigated how the RF is maintained after its formation. Previously, we showed that continued ciliary motility is required for long-term RF stability (Bearce et al. (2022b)). Here, using the temperature-sensitive *cfap298* mutant, we tested how RF integrity responded to acute reduction of motility. Embryos were raised at permissive temperature for 48 h to allow RF assembly, then shifted to restrictive temperature and analyzed 24 and 48 h post-shift (hps) (Fig 3A). Ciliary motility was significantly reduced by 24 hps and further diminished 48 hps (Fig 3B-3C and Video 17). To assess RF integrity, we imaged Sspo-GFP across five evenly spaced axial regions along the CC (Fig 3D). At 24 hps, the RF remained largely intact, especially rostrally, but showed increased flocculence caudally (Fig 3E). By 48 hps, the RF had degraded in nearly all embryos, with diffuse Sspo material accumulating within the CC (Fig 3F). We conclude that sustained ciliary motility is essential for RF stability, and that disassembly occurs rapidly once motility is impaired, especially in caudal locations. This likely reflects a requirement for cilia motility to incorporate new Sspo material to maintain fiber.

**Figure 3.**
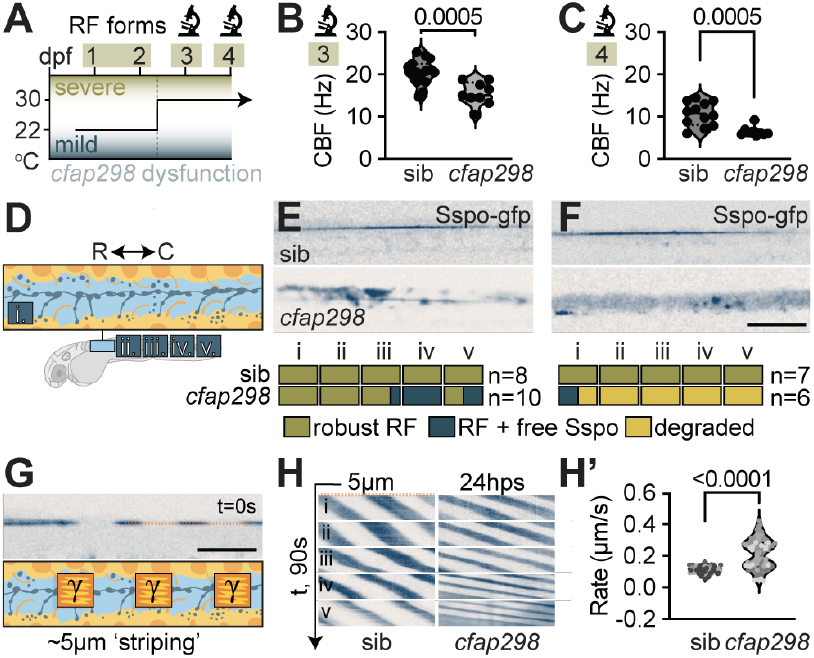
Temperature shift assays and RF translocation. **A)** Schematic of temperature-shift experiments initiated after RF assembly. The dashed line indicates the day of shift. **B-C)** Median CC CBF in *cfap298* embryos at 24 (B) and 48 (C) hps. P values are shown (unpaired t-test). **D)** Schematic of 5 imaging windows (i-v) spaced 5 somites apart. **E-F)** RF assembly and qualitative phenotyping at 24 (E) and 48 (F) hps. **G-H’)** Schematic of RF photobleaching for translocation analysis. Pixels along the dashed line were used to generate kymographs **(H)** from axial locations i-v. A steeper slope indicates faster RF translocation, quantified in (H’). P values are shown (unpaired t-test). Scale bars: 10 µm.

The RF is a dynamic structure which continuously translocates caudally, a movement thought to result from ciliary motility and the hydrodynamics within the CC. To assess translocation, we photobleached stripes into intact fibers at 24 hps and live-imaged RF movement (Fig 3G). In controls, the RF translocated caudally at 0.11 ± 0.02 µm s^*−*^1 (mean ± s.d.; Fig 3H-3H’ and Video 18). Contrary to the prediction that movement would slow under motility-impairment, translocation rates nearly doubled in *cfap298* mutants at 24 hps to 0.21 ± 0.09 µm s-1 (Fig 3H-3H’ and Video 18). This suggests that motile cilia normally contribute to a resistive effect that slows RF caudal translocation.

### FP fibrils anchor the RF and regulate tension release

The unexpected increase in RF translocation in the reduced ciliary motility condition of temperature shifted *cfap298* mutants suggested that forces act on the fiber beyond ciliary motility. Prior work proposed that the RF has elastic properties and exists under tension *in vivo* (Dendy (1909); Bellegarda et al. (2023)). To test this, we combined photobleaching, creating discrete stripes on the RF where fluorescence was absent, with acute photoablation to sever the RF (Fig 4A). Following severing, photobleached stripes contracted significantly, with a post-/pre-cut length ratio of L1/L0 = 0.55 ± 0.08 (mean ± s.d.; Fig 4A-4A’), confirming that the RF behaves as an elastic thread under tension.

**Figure 4.**
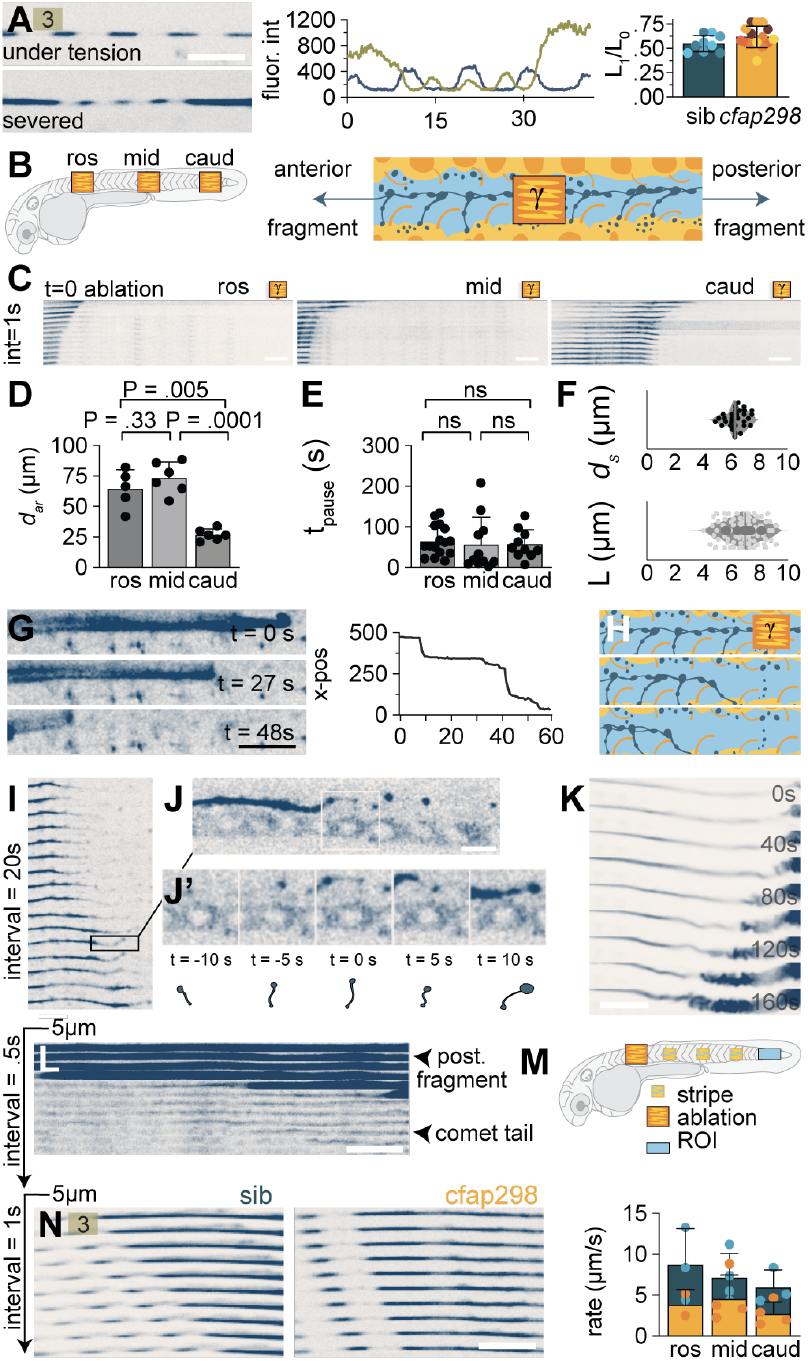
Retraction dynamics. **A)** Photobleached RF segment in 72 hpf wild type at its initial position (upper) and 180 s post-severing (lower). The striped (photobleached) segments shorten due to retraction (A’). **B)** Schematic of ablation positions where RF was severed used in C-D (left). After a cut, the RF retracts both rostrally and caudally from the cut site. **C)** Time series along the RF (thin strip centered on the RF axis) showing the initial rapid recoil of the anterior fragment after rostral, middle and caudal severing. **D)** Distance of initial anterior recoil (*d*_*ar*_) from ablations in C. **E)** Pause duration (*t*_*pause*_) between stepwise retraction events after ablations in C. **F)** Mean step size during anterior retraction (*d*_*s*_) (upper) and mean FP cell length (lower). **G)** Manual tracking of the severed end identifies discrete pauses at the site of FP fibrils (left), revealing a stepwise catch-and-release cycle (right). **H)** Schematic illustrating fibril tilt and reorientation during anterior retraction. **I)** Time series along the RF during fiber regrowth after severing. **J-J’)** Magnified view from I showing a fibril reorienting beneath the regrowing RF. Fibrils are schematized (J’, lower). **K)** The posterior fragment accumulates in front of the caudal ampulla after severing. **L)** Time series of posterior retraction showing smooth, pause-free movement toward the caudal ampulla, with comet-like Sspo trails. **M)** Schematic illustrating the positions of photobleaching (stripes) and rostral ablation for measuring the rates of RF retraction as stripes enter the caudal ROI. **N)** (Left) Time series of posterior retraction showing that stripe movement is faster in sibling controls compared to *cfap298* mutants 24 hps. (Right) Rate at rostral, middle and caudal stripes enter the caudal ampulla in sibling controls and *cfap298* mutants 24 hps. Scale bars: 10 µm.

Severing in rostral, middle or caudal locations each generated two fragments: an anterior fragment that retracted rostrally and a posterior fragment that retracted caudally (Fig 4B; Video 19; (Bellegarda et al. (2023)). Focusing first on the anterior fragment, we found that the initial recoil distance (dar) varied depending on the axial location where the RF was cut. Cuts in rostral and middle regions produced large recoils (65.5 ± 15.7 µm and 73.7 ± 13.1 µm, respectively; Fig 4C-4D), whereas caudal cuts produced smaller recoil (26.9 ± 4.7 µm; Fig 4C-D). This indicates that the RF stores pre-existing tension and that this tension is graded, diminishing caudally. This is consistent with a prior report of greater RF oscillations near the tail (Bellegarda et al. (2023)). Recoiling ends often appeared frayed (Video 20), suggesting partial rupture of lateral interactions within the fiber.

After recoil, anterior fragments continued to retract caudally at a slower speed (Fig 4C). This phase occurred in a stepwise fashion, punctuated by pauses of 60.6 ± 47.0 s (Fig 4C, 4E and Video 21). During pauses, fragments visibly caught on FP-anchored fibrils (Fig 4G and Video 22). Detachment from a fibril triggered renewed retraction, creating a catch-and-release cycle as the fiber was temporarily restrained by catching on successive fibrils (Fig 4G-H). Indeed, the mean step size (7.4 ± 2.7 µm) closely matched FP cell length (6.8 ± 1.2 µm; Fig 4F), indicating that fibrils anchored successive FP cells.

Fibril orientation changed dynamically during this process. In unperturbed embryos, fibrils tilted caudally, aligned with the direction of RF translocation. During anterior retraction, fibrils adjacent to the posterior end of the retracting anterior fragment flipped rostrally, resisting retraction before releasing and allowing the RF to snap forward (Video 22). This shows that fibrils actively reorient to resist tension rather than serving as static attachments (Fig 4H). To further assess fibril reorientation, we visualized RF regrowth, which occurred from the rostral end after severing-induced retraction (Fig 4I). In regions without intact fiber, fibrils extended dorsally from the FP and shifted slightly between rostral and caudal orientations, likely influenced by local flows (Fig 4J-4J’). However, once incorporated into the regenerating RF, fibrils consistently adopted a caudal tilt (Fig 4J-4J’ and Video 23), indicating that RF attachment imposed directional tension and aligned fibrils with the posterior forces that drive fiber translocation.

Thus, the RF is an elastic thread whose tension is constrained by FP fibrils. Severing induces rapid rostral recoil followed by gradual, stepwise retraction as fibrils catch and release the fiber (Fig 4H). These fibrils dynamically reorient to oppose recoil and then realign caudally when reengaged with the fiber. Fibril anchorage explains the increased RF translocation observed when ciliary motility was reduced by temperature shifts in *cfap298* mutants (Fig 3H-H’): impaired motility reduces fibril formation, lowering anchorage and thereby increasing caudal translocation. Axial differences in recoil likewise reflect the number of FP anchors engaged: caudal cuts left longer anterior fragments attached to more FP cells, dissipating tension across multiple anchors, whereas rostral cuts left shorter fragments with fewer anchors, permitting greater recoil.

### Posterior fragments accelerate without stepwise restraint

We next analyzed posterior fragments generated by RF severing, which retracted smoothly into the caudal ampulla and accumulated there as a coiled mass (Fig 4K and Video 24). Unlike retracting anterior fragments, this motion showed little pause or visible resistance from fibrils. We propose that this difference reflects fibril orientation: in the intact state, fibrils are tilted caudally, so anterior retraction pulls against them and produces transient resistance, whereas posterior retraction proceeds with fibril orientation and therefore encounters weaker restraint. During posterior retraction, we frequently observed comet-like trails of Sspo following the retracting fragment, likely flocculent material swept up as the fiber moved caudally through the CC (Fig 4L and Video 25). To quantify posterior retraction, we photobleached discrete stripes in rostral, middle, and caudal regions of intact fibers and then severed the RF rostral to all photobleached sites (Fig 4M). Caudal stripes arrived at the ampulla first, moving at a rate of 6.0 ± 2.1 µm s-1 (mean ± s.d.; Fig 4N and Video 26). This was followed by middle stripes (7.1 ± 3.0 µm s-1) and finally rostral stripes (8.7 ± 4.4 µm s-1; Fig 4N and Video 26). Thus, posterior fragments accelerate as they retract. This is consistent with the idea that residual anchorage to the FP decreases as the fragment approaches the caudal end, allowing progressively faster motion.

In *cfap298* mutants at 24 hps—when ciliary motility was impaired, but the RF remained intact and tensioned (Fig 4A)—posterior retraction was significantly slower (Fig 4N and Video 27). We interpret this reduction as evidence that motile cilia contribute forces and flows that promote caudal fiber movement. Supporting this, double severing in wild type generated tensionless RF segments that continued to transit caudally, albeit more slowly than singly cut fibers (Video 27), indicating that tension release alone is insufficient to drive retraction. Together, these findings revealed a striking asymmetry in RF recoil. Anterior fragments retract stepwise as anchoring fibrils transiently resist tension release, whereas posterior fragments accelerate smoothly into the tail, a process driven by stored tension but further aided by motile cilia-driven flow. This reinforces the model that fibril orientation favors posterior but resists anterior RF motion. Additionally, this suggests motile cilia play a dual role in both of these functions, promoting fibril formation and anchorage, permitting tension buildup while facilitating posterior translocation.

### The RF drives axial straightening through spatial and temporal control

Last, we investigated how spatial and temporal aspects of RF formation contribute to axial morphogenesis. In wild-type embryos, the body transitions from a ventrally curved posture at 16 hpf to a straightened axis by 24–26 hpf (Bearce and Grimes (2020)). Loss of RF formation—whether through impaired cilia motility or absence of Sspo—prevents straightening and produces the curly tail down (CTD) phenotype (Kramer-Zucker et al. (2005); Jaffe et al. (2016); Cantaut-Belarif et al. (2018)). Because the RF assembles progressively from rostral to caudal regions of the CC, we asked whether partial, region-restricted assembly would support straightening locally while failing in regions of unassembled RF.

To test this, we examined a spontaneous mutant with a curved body, *cebra*, which we mapped to a novel allele of *ruvbl2*, a gene required for ciliary motility in zebrafish (Fig 5A; (Zhao et al. (2013)). *cebra* mutants displayed two phenotypic classes: some failed to straighten entirely, resulting in severe CTD, while others showed a milder “hook-tailed” phenotype, with rostral straightening but caudal curvature (Fig 5C). Live imaging across five defined CC regions (Fig 3D) revealed that ciliary motility was significantly reduced in CTD mutants but, in hook mutants, slightly elevated rostrally and reduced caudally (Fig 5B and Video 28). RF integrity mirrored this gradient: siblings consistently formed a continuous RF along the CC, severe CTD mutants lacked an RF throughout, while hook-tailed mutants retained a continuous rostral RF but lost it caudally (Fig 5D and 5I). Thus, RF integrity correlated with regional straightening, supporting a spatially localized role for the RF in axial morphogenesis.

**Figure 5.**
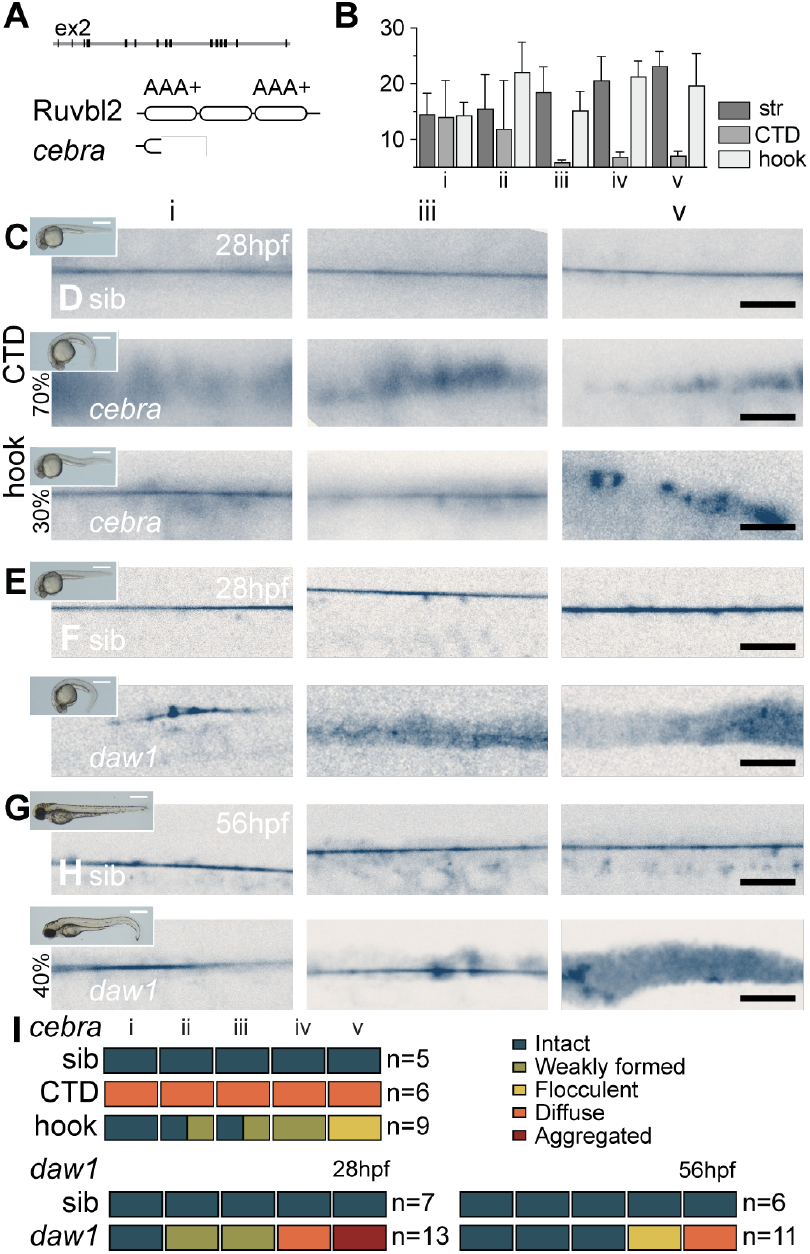
Spatial and temporal assembly. **A)** *cebra*, a spontaneous *ruvbl2* mutant, alters a splice site and is predicted to cause an early truncation. Regions that assemble into the AAA+ ATPase domain are shown. **B)** Median CBF for sibling, *cebra* CTD and *cebra* hook mutants across five axial locations (see Fig 3D). Error bars show s.d. **C)** Body axis phenotypes at 28 hpf, showing two classes of cebra mutants. **D)** Sspo-GFP imaging at the indicated axial positions at 28 hpf. RF is absent in cebra CTD mutants but present rostrally in cebra hook mutants. **E)** Body axis phenotypes in daw1 mutants at 28 hpf. **F)** Sspo-GFP imaging at the indicated axial positions at 28 hpf. daw1 mutants form only weak RF rostrally and occasionally in mid-CC regions, with absent caudal assembly. **G)** Body axis phenotypes in daw1 mutants at 56 hpf, when partial recovery of CTD has occurred. The most common phenotype (40%) is shown. **H)** Sspo-GFP imaging at the indicated axial positions at 56 hpf. Recovering mutants showed substantially greater RF assembly than at 28 hpf. **I)** Summary of Sspo-GFP imaging data from panels D, F, and H. Scale bars: 0.5 mm (C, E, G); 10 µm (D, F, H).

We next examined temporal requirements for motile cilia using *daw1* mutants (Bearce et al. (2022a)). Daw1 promotes dynein arm import into cilia (Ahmed and Mitchell (2005)), and loss of *daw1* delays CC cilia motility, with low levels of motility at 28 hpf that had significantly increased by 56 hpf (Bearce et al. (2022a)). This recovery paralleled body straightening: *daw1* mutants displayed CTD at 28 hpf but progressively straightened between 32-120 hpf (Fig 5E and 5G; (Bearce et al. (2022a))). Here, we found that RF dynamics followed this same timeline. At 28 hpf, mutants only formed a thin rostral RF, with caudal regions lacking a fiber (Fig 5F and 5H), but by 56 hpf the RF extended caudally in step with cilia motility and body axis recovery (Fig 5H and 5I). Together, these findings showed that RF function is both spatially and temporally regulated, with region-specific assembly and developmental timing playing key roles in axial morphogenesis.

## Discussion

In summary, our results reveal a multi-step process by which secreted Sspo is remodeled into a tensioned, posteriorly translocating Reissner fiber (RF) that spans the central canal (CC). Classic histological and ultrastructural studies suggested that subcommissural organ (SCO) secretions form fine filaments that condense into the RF, potentially aided by ciliary activity (Afzelius and Olsson (1957); Castenholz and Zöltzer (1980); Wislocki and Leduc (1952)). Work in zebrafish later demonstrated that cilia motility is required for RF formation, as mutations disrupting beating prevent assembly (Cantaut-Belarif et al. (2018); Rose et al. (2020); Bearce et al. (2022b); Wang et al. (2022)). Our findings build on and mechanistically extend this model, revealing that motile cilia actively remodel secreted Sspo globules into elongated fibrils that then twist and bundle into a continuous, tensioned fiber.

This process bears striking resemblance to the spinning of wool, in which loose fibers are drawn out into thin strands before being twisted together into a strong, cohesive thread. In textile spinning, twisting aligns fibers, binds them through friction and entanglement, and progressively thickens the yarn as additional material is incorporated. Likewise, we find that the RF assembles by stretching flocculent Sspo globules into elongated fibrils that twist and bundle into a fiber. As in spun wool, new material is continuously recruited in a helical fashion into an aligned, tensioned scaffold, transforming the Sspo into a structure capable of bearing mechanical load. This mechanism may help explain how, in large-bodied animals such as cattle, the RF can grow to astonishing thickness, reaching diameters greater than a third of millimeter (Estivill-Torrús et al. (1998)). Ultrastructural studies in bovine brain support this view, revealing that the RF consists of bundled fibril-like structures (Muñoz et al. (2019)). Additional proteins, including Galectin-1, have also been implicated in fiber stability, potentially by cross-linking threads (Muñoz et al. (2019)), though this remains to be tested through loss-of-function approaches.

The domain architecture of Sspo is consistent with our model of RF assembly. Its N-terminal region contains multiple von Willebrand factor (vWF) D and C domains, which in the blood-clotting protein vWF are exposed by shear-induced stretching, unmasking adhesive sites that mediate multimerization and platelet cross-linking (Okhota et al. (2020)) and this transition is favored by tethering (Wang et al. (2019), reflective of Sspo’s extrusion at the site of FP-anchored fibrils. Similarly, mechanical forces generated by motile cilia and CSF flow may stretch Sspo globules into fibrils, exposing adhesive vWF domains and thereby promoting lateral interactions during RF assembly. In addition, Sspo harbors numerous thrombospondin type 1 (TSR) repeats, known to mediate adhesion and binding to extracellular matrix proteins. Within the RF, these TSR domains may reinforce fibril bundling and enhance the mechanical integrity of the fiber. A recent study demonstrated that the C-mannosyltransferase Dpy19l1l modifies TSRs of Sspo and is essential for RF formation (Tian et al. (2025)), reinforcing the idea that biochemical modification of Sspo, not just its secretion, is a critical step in fiber assembly.

Beyond its assembly, our results show that the RF is mechanically anchored to the FP by fibrillar connections distributed along its length. These connections are not static attachments but actively respond to surrounding forces, reorienting to constrain local fiber movement and enabling tension to build along the RF. This dynamic anchoring system, together with motile cilia activity, allows the RF to withstand mechanical forces and potentially transmit mechanical information to the FP. Future investigations will be required to elucidate the nature of these RF-FP attachments.

Disrupting RF formation, either spatially or temporally, produces predictable defects in body shape, directly linking fiber assembly to axial morphogenesis. The RF can therefore be placed in a broader class of extracellular assemblies that shape morphogenesis. Across tissues and species, such assemblies transmit chemical and mechanical cues to coordinate tissue behavior (Nelson et al. (2024); Shellard and Mayor (2021); Isabella and Horne-Badovinac (2016)). The RF represents a specialized example in the vertebrate CC, where it may serve as a scaffold for posterior transport of molecular signals, or it may act a mechanical element, providing feedback to guide CC and body axis development. Indeed, mechanosensitive CSF-contacting neurons (CSF-cNs) within the CC physically contact the RF and respond with Ca2+ transients (Alibardi and Meyer-Rochow (1988); Orts-Del’Immagine et al. (2020)), raising the possibility that the fiber functions as a conduit for mechanochemical signaling. Further work will be required to determine how the hydrodynamic environment of the CC, the RF, and its cellular interactions integrate to coordinate large-scale morphogenesis of the body.

## Experimental Procedures

### Zebrafish

AB and WIK strains of *Danio rerio* (zebrafish) were used. Embryos from natural matings were incubated at 28°C unless otherwise stated. Zebrafish lines used were: *cebra*^*b1511*^ (this manuscript), *cfap298*^*tm304*^ (Jaffe et al. (2016)), *daw1*^*b1403*^ (Bearce et al. (2022a)), *sspo*^*b1446*^ (Bearce et al. (2022b)) *sspo-GFP*^*ut24*^ (Troutwine et al. (2020)). The *cebra* mutant phenotype was originally noticed in a *frem1a*^*atc280b*^ background (Eeden et al. (1996)) that had been maintained at the University of Oregon for many years; we separated the mutations with extensive outcrossing. Experiments were undertaken in accordance with research guidelines of the International Association for Assessment and Accreditation of Laboratory Animal Care and approved by the University of Oregon Institutional Animal Care and Use Committee.

### *cebra* mapping

AB strain zebrafish harboring the mutation were outcrossed to WIK strain zebrafish, and heterozygous F1 progeny were subsequently incrossed. Pools of phenotypically mutant and wild-type embryos were sorted, and total RNA and DNA was extracted at 28 hpf from 10 non-phenotypic siblings and 10 *cebra* mutants. Both bulk RNA-sequencing and wholegenome sequencing were performed, as described below. BAM files generated from RNA-sequencing data were used as input for the Mutation Mapping Analysis Pipeline for Pooled RNA-seq (MMAPR; (Hill et al. (2013))) to identify the candidate genomic region harboring the *cebra* mutation. The mutation itself was identified from whole-genome sequencing data and confirmed by Sanger sequencing (Genewiz).

### Bulk RNA-sequencing and differential gene expression analysis

Total RNA extraction was performed using the DirectZol RNA kit following manufacturer’s instructions with 10 whole embryos or trunks per clutch for n = 5 control and n = 5 mutant clutches at 28 hours post fertilization. 250 ng of RNA was brought to 50uL in Nuclease Free Water (NEB). Ribosomal RNA was removed with NEB Poly-A Selection Module and eluted in 15uL Nuclease Free Water. Depleted mRNA was used as input for Watchmaker RNA Library prep kit following the instruction manual. Finished libraries were quantified by qPCR and pooled equimolar before sequencing on Illumina NextSeq 2000. Trimming of adaptors and low-quality reads was performed using Cutadapt in R (Martin (2011)). Alignment, quantification, and analysis of RNA sequencing data was performed using the Rsubread package (Liao et al. (2019)). Additional quality control and quantification of Short Reads was done using QuasR to assess alignment (Gaidatzis et al. (2014)). Reads were aligned to a comprehensive transcript annotation for zebrafish (Lawson et al. (2020)) and converted to BAM files. Read counts and differential gene expression analysis were performed using the DESeq2 package (Love et al. (2014)). Genes with an adjusted p value greater than 0.05 were removed before analysis, resulting in only significant differentially expressed genes. Genes with an absolute value log2-fold change greater than 1 were considered to be significant for differential gene expression. Gene ontology (GO) enrichment analysis was performed to further analyze significantly up-regulated and down-regulated genes with ShinyGO v.0.76.3 using all available gene datasets for pathway databases

### Whole genome sequencing

Whole genome sequencing was used to determine the mutation in cebra mutants. Phenol chloroform extractions were performed on wild type and mutant embryos and libraries were prepared using the FS DNA Library Prep Kit for Illumina sequencing (NEB, E7805). DNA was digested into 150 bp fragments, and paired-end sequencing was performed using a Nextseq 2000 Sequencing System. Trimming of adaptors and low-quality reads was performed by Cutadapt (Martin (2011)). Illumina short-read sequences were then aligned to the GRCz11 reference sequence of chromosome 5 using Rsubread (Liao et al. (2019)). Aligned reads in BAM format were analyzed in IGV (version 2.16.2). The mutation was subsequently confirmed using Sanger sequencing (GeneWiz).

### CRISPant generation

For mosaic mutagenesis, we routinely used four guide RNAs per targeted gene (Wu et al. (2018)). First, oligos were pooled at 10 µM then annealed and extended with 10 µM of bottom strand oligo 2 (Table 1) using Phusion HF PCR Mastermix (New England Biolabs, M05030) in a 50 µl reaction. The reaction was performed under the following thermal cycling conditions: 98°C for 2min, 50°C for 10 min, and 72°C for 10 min. Assembled oligos were purified using a DNA Clean & Concentrator kit (Zymo Research, D4013) and transcribed using a HiScribe T7 High Yield RNA Synthesis kit (New England Biolabs, E2040). The synthesized RNA was treated with 2 µL of TURBO DNase (Thermo Fisher Scientific, AM2238) for 15 min at 37°C, then purified using an RNA Clean & Concentrator kit (Zymo Research, R1013). Purified guide RNAs were aliquoted and stored at −80°C. For mutagenesis, unless otherwise stated, 1000 pg of guide RNA alongside 1600 pg/nl Cas9 (Integrated DNA Technologies, 1081058) were injected into the yolk of embryos at the 1-cell stage or the 16-cell stage.

### mRNA synthesis and expression

An *eGFP-Ruvbl2* plasmid (a gift from Zhaoxia Sun’s lab, Yale University) was linearized using NotI-HF (New England Biolabs, D4013). The purified linearized plasmid served as a template for in vitro mRNA synthesis using the mMessage mMachine SP6 Transcription kit (Thermo Fisher Scientific, AM1340). Synthesized mRNA was purified with the MEGAclear Transcription Clean-Up kit (Thermo Fisher Scientific, AM1908) and stored at −80°C. For *cebra* rescue experiments, 160 pg of mRNA was injected directly into 1-cell stage embryos.

### Immunofluorescence

Embryos were dechorionated, euthanized and then fixed in 4% paraformaldehyde (PFA) in phosphate buffered saline (PBS) overnight. Samples were then permeabilized by rocking for 90 min at room temperature in PBS + 0.1% Tween-20 (PBST), which was changed every 30 min. Samples were blocked for 2 h in 5% normal sheep serum (NSS) and 1% dimethyl sulfoxide (DMSO) in PBST at room temperature, followed by incubation with primary antibody in PBST containing 1% NSS and 1% DMSO at 4°C for 16 h. The following primary antibodies were used (1:500 dilution unless otherwise noted): anti-acetylated α-tubulin (mouse monoclonal, Sigma-Aldrich, T6793); anti-GFP (chicken monoclonal, Abcam, ab13970); anti-mCherry (chicken polyclonal, Abcam, ab205402); and anti-γ-tubulin (rabbit polyclonal, Sigma-Aldrich, T5192). Samples were washed 5 times (30 min each) in 1% NSS, 1% DMSO and 0.1 M NaCl in PBST, then incubated in secondary antibodies at 4°C in the dark. The following secondary antibodies were used: goat anti-rabbit conjugated with Alexa Fluor 546 (1:500 dilution, Thermo Fisher Scientific, A11035) and goat anti-mouse conjugated with Alexa Fluor 488 or 647 (1:500 dilution, Thermo Fisher Scientific, A11001 or A21240). Additional stains included Hoechst (1:2000 dilution, Sigma-Aldrich, 94403) and Alexa Fluor 647 Phalloidin (1:100 dilution, Thermo Fisher Scientific, A22287). Subsequently, samples were washed 5 times for 30 min per wash in 1% NSS, 1% DMSO and 0.1 M NaCl in PBST. For mounting, embryos were dissected to remove the yolk and flatten the sample, then laterally side-mounted in ProLongTM Gold Antifade Mountant (Thermo Fisher Scientific, P36930) under 1.5 coverslips. Samples were imaged using a Zeiss

LSM 880 inverted microscope with a 40x water or 63x 1.4 numerical aperture oil-immersion objective. Laser scanning confocal microscopy was performed using PMT and GaAsP-PMT detectors, with pinhole size set to 1 Airy unit. Gain settings ranged from 200-300 (GaAsP-PMT) and 550-700 (PMT). The x/y pixel size was 0.0852 µm. Images were deconvoluted using Huygens Professional, then processed and quantified in ImageJ. Cilia lengths were measured manually by tracing cilia using the Freehand Line tool in ImageJ.

### Imaging and quantitation of cilia motility

For central canal imaging, embryos between 25 and 72 hpf were manually dechorionated and anesthetized with tricaine. Embryos were laterally mounted on #1.5 coverslip-bottomed MatTek chambers by embedding them in 0.5% low-melt agarose (Thermo Fisher Scientific, BP165-25) containing tricaine, prepared in embryo medium. Imaging was performed using a Nikon Ti2 inverted microscope equipped with a Plan Apo VC 60x WI DIC 1.2 numerical aperture objective and a pco.edge sCMOS camera. Time series differential interference contrast (DIC) images (512 x 512 pixels) were acquired at 250 frames per second for 4 seconds (1000 frames total) with a pixel size of 0.11 *µ* m. Images were processed and analyzed using ImageJ. Time course data were rotated and cropped to isolate the central canal. A moving average of 55 frames were subtracted from each frame using the Subtract Moving Average function (Stowers Institute ImageJ Plugins *→* jayunruh *→* Detrend *→* Stack subtract moving average). A Gaussian blur of .8 was applied to all frames. For manual analysis, line profiles were drawn through motile cilia in FiJI (Schindelin et al. (2012)), and kymographs were generated using the Kyomograph Builder plugin. For automated analysis of motile cilia frequencies, Fourier transform-based analysis was performed using the FreQ plugin (Jeong et al. (2022)). to extract oscillations in pixel intensity associated with ciliary movement from rotated, cropped, and background-subtracted DIC images. The following parameters were used: Recording frequency — 250 Hz; Sliding window to smooth power spectrum — 0.0; Remove background from power spectra — Yes; Percent of lowest-power pixels — 2.0; Fold SD — 1.5; Extract signal regions local SD filter: Yes, 3.0 (increase range = yes); Min accepted frequency — 5 Hz; Max accepted frequency — 75 Hz.

### Quantitation of curly tail down

Larval body angle quantitation was performed as described (Bearce et al. (2022a)). Briefly, lateral images of zebrafish larvae at 1, 3 and 6 dpf were captured using a Leica S9i stereomicroscope with an integrated 10-megapixel camera. Body angles were defined as the angle formed between straight lines: one extending from the anterior to the posterior of the yolk stalk and the other extending from the posterior yolk stalk to the tip of the tail. Measurements were made in ImageJ, and statistical significance was assessed using a repeated-measures twoway analysis of variance (ANOVA).

### Live imaging of Sspo-GFP

To assess Sspo-GFP assembly and translocation, embryos and larvae (16-72 hpf) were laterally mounted for live imaging as described in the Imaging and Quantitation of Cilia Motility section. Imaging was performed using a Nikon Ti2 inverted microscope equipped with a Yokogawa W1 spinning disc, a Plan Apo VC 60x WI 1.2 NA objective, and a pco.edge sCMOS camera. Time series images (512 x 512 pixels) were acquired with a 500 ms exposure every 2 seconds for 1 minute (30 frames total). The original pixel size was 0.11 µm, with 2x2 binning applied for early-stage imaging to enhance weak signals. Qualitative descriptions of Sspo-GFP assembly were grouped into the following categories: Intact RF — a clear fiber is visible with low background noise; weakly formed RF — a partially visible fiber with high background noise and flocculent particles; flocculent Sspo — small particles accumulating into larger conglomerates; diffuse Sspo — a fine mist-like accumulation of particles is visible, with little aggregation into larger structures; aggregated Sspo — high-intensity clumps of material are collecting on the floor plate with minimal vortical movement. Photoablation. A Nikon Ti2 inverted microscope equipped with a Plan Apo VC 60x WI DIC 1.2 NA objective and pco.edge sCMOS camera was used to capture 1024 x 1024 images for 200 seconds. A fluorescence recovery after photobleaching (FRAP) module was used for photoablation, which was achieved with manual triggering of a 100 mW 405 nm laser with a 10 µm pulse diameter and 100 ms dwell time to sever the RF, for a total power delivery of 4 mW. Photobleaching was achieved at 10%.

## Acknowledgments

We thank Judy Peirce, Tim Mason and the Aquatics Facility for zebrafish husbandry, Adam Fries and the Genomics and Cell Characterization Core (GC3) Biological Imaging Facility, both at the University of Oregon. We thank Ryan S. Gray, Zhaoxia Sun and Jonathon T. Hill for sharing resources, Catherine Wilson for advice and Amy Robbins for first noticing axial curvature in the *frem1a* mutant background. This research was supported by the NIH (R35GM142949 to D.T.G., F32AR078002 to E.A.B., F31HD105435 and T32HD007348 to Z.H.I. and T32GM149387 to S.G.B.).

## Notes

### Competing Interest Statement

The authors have declared no competing interest.

